# Accurate inference of the full base-pairing structure of RNA by deep mutational scanning and covariation-induced deviation of activity

**DOI:** 10.1101/677310

**Authors:** Zhe Zhang, Peng Xiong, Tongchuan Zhang, Junfeng Wang, Jian Zhan, Yaoqi Zhou

**Affiliations:** High Magnetic Field Laboratory, Chinese Academy of Sciences, 350 Shushanhu Road, Hefei, Anhui, 230031, China; University of Chinese Academy of Sciences, Beijing, 101408, China; Institute for Glycomics, Griffith University, Parklands Drive, Southport, QLD 4222, Australia; School of Information and Communication Technology, Griffith University, Parklands Drive, Southport, QLD 4222, Australia

## Abstract

Despite the transcription of noncoding RNAs in 75% of the human genome and their roles in many diseases include cancer, we know very little about them due to lack of structural clues. The centerpiece of the structural clues is the full RNA base-pairing structure of secondary and tertiary contacts that can be precisely obtained only from costly and inefficient 3D structure determination. Here, we performed deep mutational scanning of self-cleaving CPEB3 ribozyme by error-prone PCR and showed that a library of <5×10^4^ single-to-triple mutants is sufficient to infer all 26 including nonhelical and noncanonical base pairs at the precision of a single false positive. The accurate inference, further confirmed by a twister ribozyme, is resulted from covariation analysis by utilizing both functional and nonfunctional variants for unsupervised learning, followed by restrained optimization. The result highlights the usefulness of deep mutational scanning for high-accuracy structural inference.

## INTRODUCTION

The full base-pairing structure of RNA is resulted from the interplay of secondary and tertiary interactions and serves as a preformed frame for final folding of tertiary structure. As such, it plays a prominent role in the versatility of RNA structure and function (Caprara and Nilsen, 2000; Cech and Steitz, 2014). To date, full base pairing structures at the single base-pair resolution can only be obtained from high-resolution RNA structures determined by X-ray crystallography, nuclear magnetic resonance or cryogenic electron microscopy. However, these traditional techniques solved only 4,112 RNA structures as of November 4, 2018 (or 3% of all structures in protein databank (Rose et al., 2015)) due to their requirement of nearly static structures that most RNAs do not have. Given that proteins are vastly outnumbered by noncoding RNAs (Djebali et al., 2012) and the majority of these RNAs have unknown structures and functions, alternative methods for accurate determination of RNA base-pairing structures are urgently needed.

The most economical method for locating base pairs would be a computational prediction if its accuracy could be assured. Such a computational approach is usually referred to as RNA secondary structure prediction although many base pairs are associated with tertiary interactions (Nowakowski and Tinoco, 1997). These tertiary contacts include noncanonical (non-Watson-Crick), non-nested (pseudoknot), and lone (single tertiary base pair not associated with a helix) base pairs. Despite the first secondary structure prediction method was developed nearly 50 years ago (Tinoco et al., 1971) and many advances have been made since (Seetin and Mathews, 2012; Xu and Chen, 2015), the problem remains unsolved: only 70% accuracy for predicted base pairs and 38% accuracy for secondary structure topology according to the most recent evaluation (Zhao et al., 2018). In particular, lone and noncanonical base pairs are commonly ignored in secondary structure prediction, perhaps because they are not considered as a part of the secondary structure (Nowakowski and Tinoco, 1997).

Large improvement in secondary structure prediction can be achieved (Rice et al., 2014; Somarowthu, 2016) when computational algorithms are integrated with experimental restraints. These experimental results can be generated from structural probes such as enzymes, chemicals, hydroxyl radical, cross-linking, and mass spectrometry (Harris and Christian, 2009; Meng and Limbach, 2006; Sachsenmaier et al., 2014), in combination with next-generation sequencing (Bevilacqua et al., 2016). However, most of these experimental techniques measure one-dimensional reactivity profiles (Lu and Chang, 2016) and rely on computational approaches to infer two bases paired. As a result, they suffer the same limitations common for all existing computational methods: poor accuracy in locating lone, non-nested, and noncanonical base pairs (Miao and Westhof, 2017; Rice et al., 2014; Somarowthu, 2016).

Recently, more direct identification of base pairs can be made by multidimensional chemical mapping methods. Examples are mutate-and-map (M2 (Kladwang et al., 2011)), SHAPE and mutational profiling (SHAPE-MaP (Siegfried et al., 2014)), RNA interaction groups by mutational profiling (RING-MaP (Homan et al., 2014)), multiplexed ·OH cleavage analysis (MOHCA(Das et al., 2008)), and modify-cross-link-map (MXM (Tian and Das, 2016)). While these multidimensional chemical mapping methods can detect RNA base pairs at the helix level (>2 base pairs)(Cheng et al., 2017), they are not yet sensitive enough to detect all base pairs individually, lone and two contiguous base pairs, in particular. This is in part because the assumption of a localized response upon perturbation is not always true (Tian and Das, 2016). Moreover, these experiments are labor intensive and require sophisticated computationally-intensive algorithms for data analysis (Miao et al., 2017; Tian and Das, 2016).

Another powerful technique is deep mutational scanning through function selection in combination with high-throughput sequencing (Auyeung et al., 2013; Buenrostro et al., 2014; Kobori and Yokobayashi, 2016; Li et al., 2016; Pitt and Ferre-D’Amare, 2010; Puchta et al., 2016; Smyth et al., 2015; Tome et al., 2014). However, current analysis of deep mutation data has been limited to mapping functional fitness landscape (Kobori and Yokobayashi, 2016; Li et al., 2016; Pitt and Ferre-D’Amare, 2010; Puchta et al., 2016; Tome et al., 2014) or locating helices via simple covariation analysis (Auyeung et al., 2013; Buenrostro et al., 2014; Smyth et al., 2015; Tome et al., 2014). Separately, sophisticated statistical covariation analysis such as evolutionary coupling, or direct coupling has been successfully applied to naturally occurring mutant variants within a large RNA family (De Leonardis et al., 2015; Weinreb et al., 2016). However, this analysis requires a large number of known homologous sequences within the same functional family, of which the majority of noncoding RNAs do not have. Moreover, the accuracy of such analysis strongly depends on the quality and quantity of homologous sequences.

Unlike naturally occurring homologous sequences, the quality and quantity of functional mutants can be somewhat controlled in deep mutational scanning. Thus, it is highly desirable if deep mutation data alone can be used to reliably infer complex secondary and tertiary base pairing structures including lone base pairs not detectable by current chemical probing techniques. We demonstrate the feasibility by employing self-cleaving ribozymes that catalyze their own cleavage. These ribozymes are particularly suitable for deep mutational scanning (Kobori and Yokobayashi, 2016) because the functional activity of a mutant can be calculated by simply counting the number of cleaved and uncleaved reads of that mutant in high-throughput sequencing data. More importantly, self-cleaving ribozymes have complex base-pairing structures, each of which has its own interesting mechanistic difference (Jimenez et al., 2015b; Liu et al., 2017; Ren et al., 2017). Thus, they offer an ideal testing ground for inferring different base pairing topologies from deep mutational scanning.

Here, we performed deep mutational scanning of the self-cleaving 81-nucleotide CPEB3 ribozyme located at the intron region of the human gene of cytoplasmic polyadenylation element–binding protein 3 (CPEB3). Although the tertiary structure of CPEB3 ribozyme remains to be solved, it will fold into a hepatitis delta virus-like base pair pattern confirmed by mutational studies (Salehi-Ashtiani et al., 2006). This ribozyme was chosen for its complex base-pairing structure. One pseudoknot is non-helical, made of one lone canonical Watson-Crick (WC) pair and one noncanonical pair and the other is capped by two noncanonical base pairs, all associated with tertiary interactions (Figure 1A). Moreover, this ribozyme is of biological significance because a single nucleotide polymorphism was found to affect CPEB3 ribozyme activity with a difference in episodic memory (Vogler et al., 2009). We established a method that analyzes covariation-induced deviation of activity (CODA) by using Support Vector Regression (SVR) to establish a mutation-independent model and a naïve Bayes classifier to separate bases paired from unpaired. This unsupervised CODA analysis improves the signal-to-noise ratio by employing both functional and nonfunctional variants. As a result, it increases the area under the precision-recall curve by >35% over classical mutual information (MI) (Dunn et al., 2008) and state-of-the-art evolutionary or direct coupling analysis (DCA) (Feinauer et al., 2014; Morcos et al., 2011) that rely on functional variants only. Moreover, incorporating Monte-Carlo simulated annealing with a simple energetic model and a CODA scoring term further improves the coverage of the regions under-sampled by mutations. Accurate determination of secondary and tertiary base pairs is further confirmed by CODA-based analysis of the available deep mutation data of the downstream portion of the cleavage site of a twister ribozyme (a 48-nucleotide fragment of 54 nucleotide *Oryza sativa* Osa-1-4 ribozyme sequence) (Kobori and Yokobayashi, 2016).

**Figure 1.**
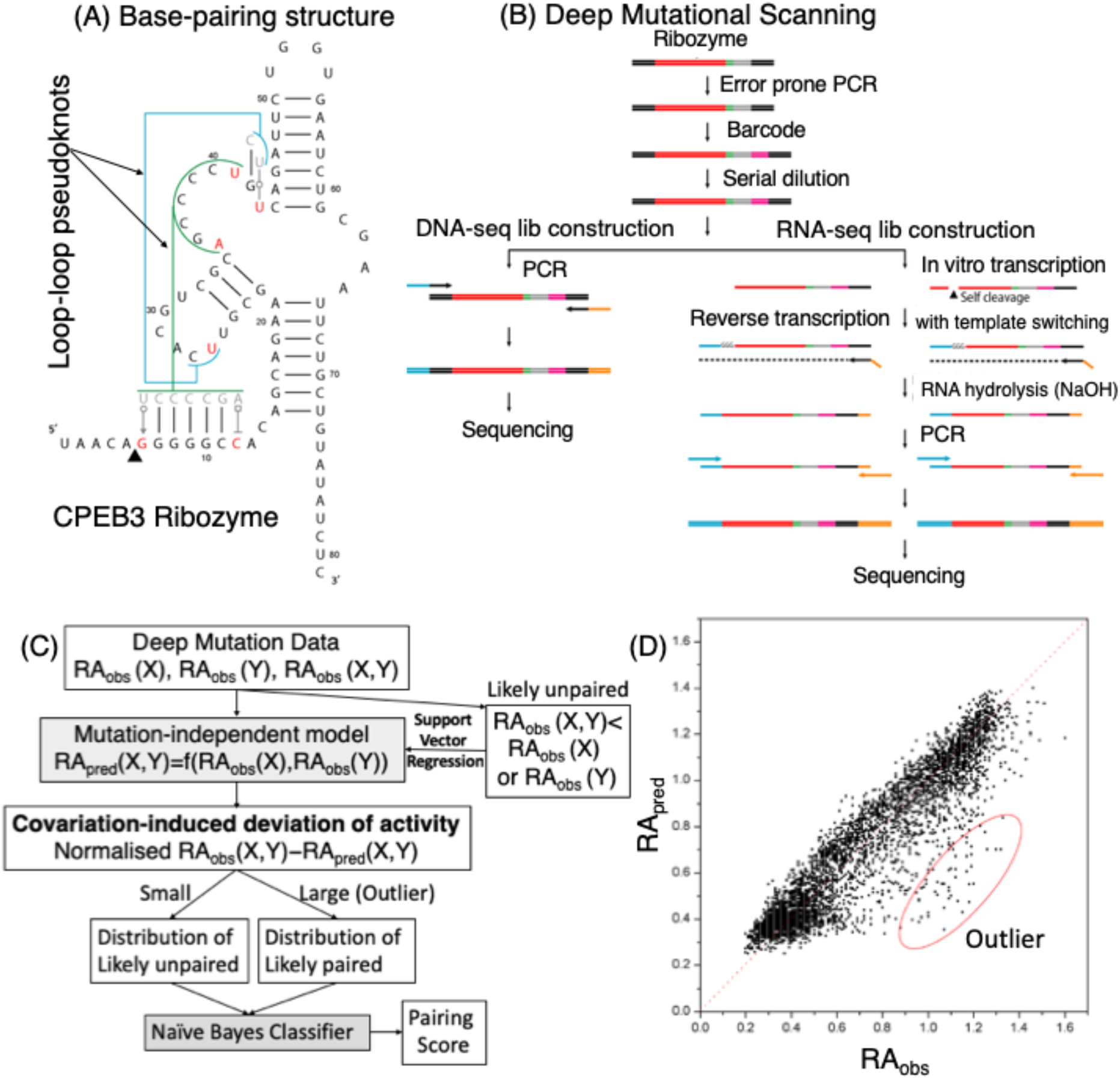
(A) The native base-pairing structure of the 81-nucleotide CPEB3 ribozyme contains three canonical helical regions. In addition, it has one pseudoknot capped by two non-Watson-Crick (WC) base pairs (G6U41 and C12A35) and another pseudoknot made of a lone WC pair and one non-WC pair (U26U43). (B) The deep mutational sequencing of CPEB3 ribozyme starts from random mutagenesis via error-prone PCR, followed by barcoding so that the whole ribozyme sequences in DNA-seq can be mapped onto self-cleaving reaction products from the RNA-seq library. (C) Analyzing deep mutation data by covariation-induced deviation of activity. This is done by constructing a mutation-independent model for those likely unpaired bases (the activity of double mutant is lower than one of the single mutants. i.e., no covariation-induced recovery of activity) and building a naïve Bayes classifier for base pairs according to distributions of unlikely paired bases and likely paired bases (outliers of the mutation-independent model). (D) The activities of double mutants observed from sequencing versus those predicted by the mutation-independent model. Outliers for likely base pairs are indicated by the red circle.

## RESULTS

### Deep mutational scanning experiment

The deep mutational scanning experiment of CPEB3 ribozyme was carried out by random mutations generated from error-prone PCR (epPCR) as shown in Figure 1B with mutation rates varied from 1.73% to 5.62% in three different batches so as to maximize the coverage of double and triple mutations. The resulting mutation library was barcoded and separated for DNA sequencing (DNA-seq) and *in vitro* transcription to RNAs, respectively. Transcribed RNAs self-cleaved during the transcription process. These self-cleaved and non-cleaved RNAs were then reverse-transcribed back to DNA for sequencing (RNA-seq). DNA-seq was used to obtain the full sequences of mutation variants that can be mapped to the reaction products from RNA-seq according to barcodes. This mapping was necessary to calculate the relative activity of each variant based on the fraction of cleaved fragments of the variant from RNA-seq, relative to that of the wild-type sequence. Comparing to multi-dimensional chemical probing and 3D structure determination, these experimental steps are relatively simple and straightforward. Combining three batches of experiments led to the deep mutation data of CPEB3 ribozyme with a total of 243 single, 11,968 double, 36,214 triple and 62,992 other (>3) mutants, along with measured enzymatic activities. This represents 100% coverage of single mutations but only 41.0% coverage of all possible double mutations. The low coverage of double mutations is in part because error-prone PCR tends to have fewer mutations in the GC than in the AT region (Cadwell and Joyce, 1992; Keohavong and Thilly, 1989). To expand the coverage of double mutations, we approximated all triple as double mutations (see Methods) and improved the coverage to 61.3% of possible double mutants with 86.9% for AT and 31.0% for GC pairs. The low mutation coverage for GC pairs and the small library (<50,000 single-to-triple mutation variants) makes it challenging for covariation analysis.

### Covariation-induced deviation of activity (CODA)

The deep mutation data obtained above was employed to search for the base pairs whose covariation (co-mutation) would lead to the recovery of cleavage activity, disrupted by single mutations. To build a sensitive method, we first employed support vector regression to establish a mutation-independent model in which the relative activity of a double mutant is predicted by the activities of two corresponding single mutants (Figure 1C). This model was built by those double-mutation variants with activity lower than one of the single mutants (i.e. lacks activity recovery from single mutations, thus, likely unpaired bases). Then, the deviation of the observed activity from the predicted activity measures the strength of covariation, with outliers associated with potential base pairs (Figure 1D). The distribution of these covariation-induced deviations of activity (CODA) can be deconvoluted into two separate distributions for independent double mutations centered at CODA=0 and outlier double mutations for likely base pairs, respectively (Figure 1C, see Methods). A naïve Bayes classifier was obtained to estimate the probability of two bases paired. Both the SVR model and the Bayes classifier were obtained from the deep mutation data alone without making any explicit/implicit assumptions about which two bases are paired or not paired (i.e. unsupervised). In other words, it is a predictive model applicable to RNAs with unknown base-pairing structures.

### Performance

Table 1 compares the performance of CODA on inferring true base pairs from the deep mutation data of CPEB3 ribozyme with those of two covariation analysis tools MI and DCA commonly used for obtaining contact information from functional homologous sequences (see Methods). DCA has essentially the same performance as MI with the Matthews correlation coefficient (MCC) value of 0.56, the sensitivity of 35% and the precision of 90%. Note that MCC is 1 for perfect prediction and 0 for random prediction. Although their overall performance is lower than secondary structure predictors (either RNAfold (MCC=0.66) or IPknot (MCC=0.71)), RNAfold can only predict nested helical stems. IPknot can predict the pseudoknot with the long contiguous region of stacked base pairs but not the short, nonhelical one. By comparison, MI, DCA, and CODA can all capture some nested along with some non-nested base pairs from both pseudoknots as shown in Figure 2A. CODA makes a large improvement over MI and DCA with 42% and 35% increase, respectively, in the area under the precision-recall curve (AUC_PR, 1 for perfect prediction), 13% and 11% increase, respectively, in MCC. Systematic improvement of CODA over MI and DCA is clearly demonstrated in the precision-recall curve (Figure 2B). MI and DCA yield 9 correct base pairs and 1 false positive prediction, while CODA produces 11 correct base pairs at 100% precision. It is particularly encouraging to note that all covariation analysis (MI, DCA, and CODA) can capture one (UU) or two (UU and GU) out of three noncanonical pairs. Neither RNAfold nor IPknot can predict any noncanonical pairs as these base pairs are commonly ignored in secondary structure prediction.

**Table 1.** Performance of mutual information (MI), direct coupling analysis (DCA), covariation-induced deviation of activity (CODA, this work), Monte Carlo simulated annealing (MC), and CODA+MC for inferring base pairs from deep mutation data in term of area under the precision-recall curve (AUC_PR), Matthews correlation coefficient (MCC), sensitivity, and precision for CPEB3 and twister ribozymes. Results for RNA secondary structure predictors (RNAfold and IPknot) are also shown for comparison.

| Method | AUC_PR | MCC | Sensitivity | Precision |
| --- | --- | --- | --- | --- |
| CPEB3 (81 bases, 26 base pairs) |  |  |  |  |
| RNAfold | - | 0.66 | 0.65 | 0.68 |
| IPknot | - | 0.71 | 0.73 | 0.70 |
| MI | 0.38 | 0.56 | 0.35 | 0.9 |
| DCA | 0.40 | 0.56 | 0.35 | 0.9 |
| CODA | 0.54 | 0.62 | 0.39 | 1 |
| MC | 0.54 | 0.56 | 0.85 | 0.37 |
| CODA+MC | 0.998 | 0.98 | 1.0 | 0.96 |
| Twister Fragment (48 bases, 17 base pairs) |  |  |  |  |
| RNAfold | - | 0.22 (0.68<sup>a</sup>) | 0.18 (0.47<sup>a</sup>) | 0.30 (1<sup>a</sup>) |
| IPknot | - | 0.0 (0.68<sup>a</sup>) | 0.0 (0.47<sup>a</sup>) | 0.0(1<sup>a</sup>) |
| MI | 0.47 | 0.56 | 0.53 | 0.60 |
| DCA | 0.06 | 0.19 | 0.53 | 0.09 |
| CODA | 0.71 | 0.79 | 0.76 | 0.81 |
| MC | 0.60 | 0.68 | 0.65 | 0.73 |
| CODA+MC | 0.91 | 0.91 | 0.82 | 1 |

<sup>a</sup>Folded using the whole sequence (54 bases), rather than the 48-base fragment.

**Figure 2.**
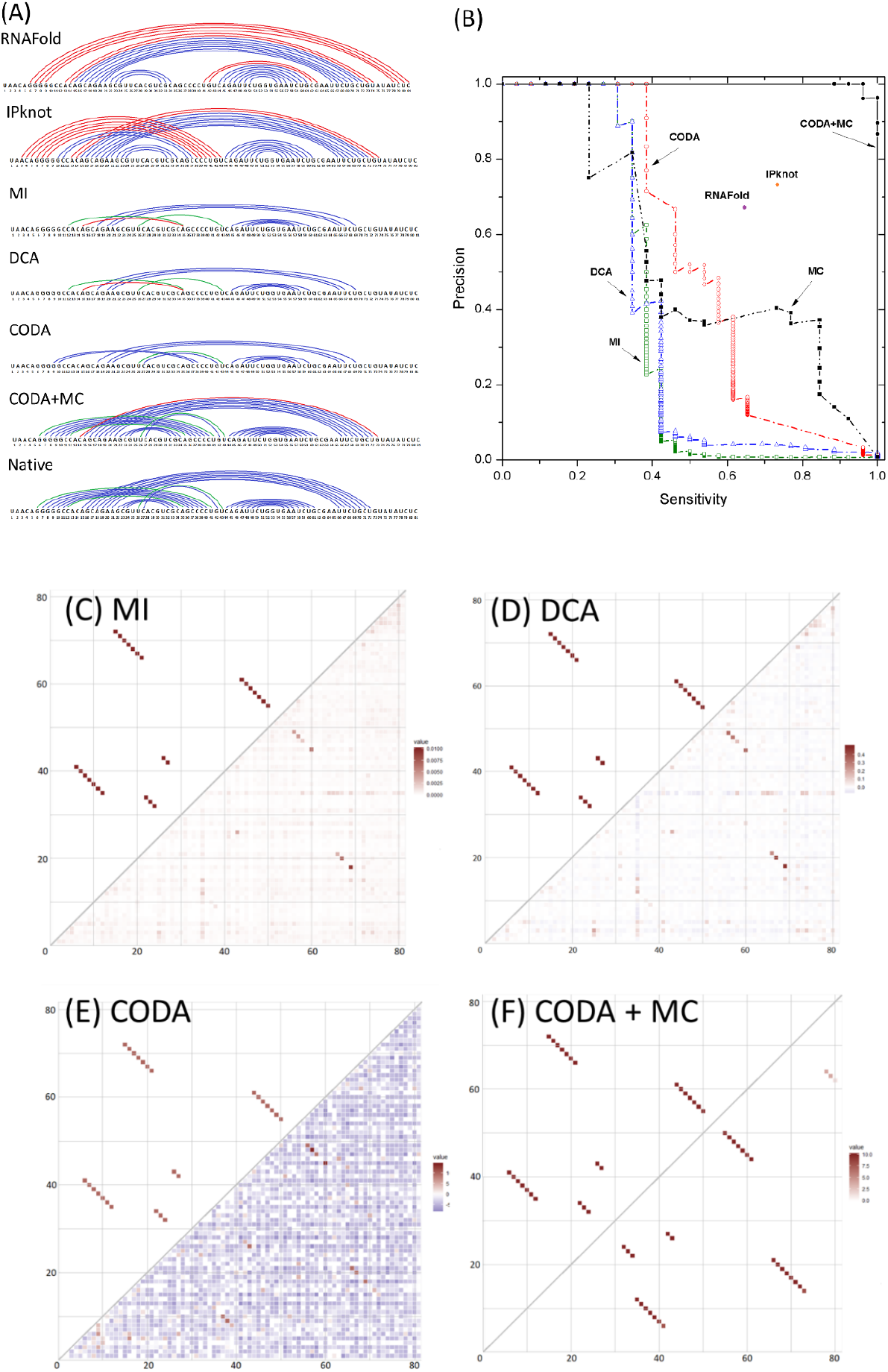
(A) The native base pairs of the CPEB3 ribozyme along with base pairs predicted by secondary structure predictors RNAfold and IPknot and inferred from its deep mutation data by mutual information (MI), direct coupling analysis (DCA), covariation-induced deviation of activity (CODA), and CODA integrated with a Monte Carlo simulated annealing (CODA+MC). Native Watson-Crick and non-Watson-Crick base pairs are shown in blue and green, respectively. False positive predictions are shown in red. (B) Precision (fraction of correct base pairs in predicted base pairs) versus sensitivity (coverage of known base pairs) by MI, DCA, CODA, MC, and CODA+MC, using the deep mutation data from CPEB3 ribozyme. Results from secondary structure predictor (RNAfold and IPknot) are also shown as points. (C) The comparison between native base-pairing map (upper triangle) of CPEB3 ribozyme and the map inferred from CPEB3 ribozyme deep mutation data (lower triangle) by MI. (D), (E), and (F) are the same as (C) but for DCA (D), CODA(E) and CODA+MC (F), respectively.

What makes CODA more sensitive in detecting base pairs? Figures 2C-E show the density plot for base-pairing maps given by MI, DCA, and CODA, respectively. The predicted scores for all base pairs by CODA have the widest distribution, suggesting the large contrast between the scores for bases paired and those for bases unpaired. This is confirmed by Supplementary Figure S1, which compares the score distributions of bases unpaired, paired with double mutations fully covered, and without fully covered (lack of statistics). These three states are well separated in the distribution of *Ps* scores in CODA, but neither in MI nor in DCA. This confirms the significant improvement of the signal-to-noise ratio by CODA over MI and DCA, due to the fact that CODA employs both functional and non-functional sequences whereas MI and DCA rely on functional sequences only.

However, the helix regions derived from deep mutation data are shorter than the native ones (Figure 2A). Undetected base pairs usually occur at the edge of the region with contiguous base pairs because the mutation at the edge of a stem affects less on structural stability and thus leads a weaker covariation signal. Moreover, the mutant library is small and biases toward AT pairs because error-prone PCR tends to have fewer mutations in the GC than in the AT region (Cadwell and Joyce, 1992; Keohavong and Thilly, 1989). The low mutation coverage leads to low base-pair coverage (or low sensitivity of MI, DCA, and CODA).

To remedy the problem of underestimating the length of contiguous base-paired regions, we introduced a simple base-pair predictor by using experimentally measured base-pair and stacking energies (Xia et al., 1998) and a restraint that one base cannot be paired with more than a single base. We employed a single adjustable parameter to penalize lone pair and non-AU, GC, or wobble GU pairs by setting it to 3 kcal/mol so that the predictor does not produce any lone and base pairs beyond AU, GC, and GU in base-pair prediction by Monte Carlo (MC) simulated annealing (results are not sensitive to the value between 2-5 kcal/mol). As shown in Table 1, MC alone can yield a reasonable performance (although not as accurate as RNAfold in term of the MCC value). Then, we incorporated the pairing score (*Ps*) from CODA as an additional energetic term with a weight factor of 1 because the magnitude of the CODA term happens to be similar to experimental base-pairing energies used in MC. We performed MC simulated annealing with random initial seeds 100 times in order to obtain contact probabilities. Table 1 shows that incorporating CODA to MC increases the number of correctly predicted base pairs from 11 to all 26 base pairs with a single false positive as shown in Figure 2A. A near perfect precision-recall curve by CODA+MC (Figure 2B) highlights the power of combining a base-pair predictor with CODA analysis. We noted that the falsely predicted pair is a GC pair (C14G73) at the end of a helix with 7 base pairs.

### Twister Ribozyme

To confirm the improvement of CODA analysis over MI and DCA and high-accuracy inference of base-pairing structure by CODA+MC, we obtained the raw deep mutation data of a twister ribozyme from Kobori and Yokobayashi (Kobori and Yokobayashi, 2016). The ribozyme can self-cleave into two RNA fragments of 6 and 48 bases long. The ribozyme mutant library is in the C-terminal 48-nucleotide fragment only. Thus, we will focus on this fragment whose base-pairing structure (Liu et al., 2014) is shown in Figure 3 (pdb ID 4OJI). The base pair pattern contains three noncanonical pairs (A2G39, A22A40, and Hoogsteen base pair U18A22), one lone pair (C9G13) as well as two long-distance loop-loop pseudoknots (or kissing interactions) that were considered as tertiary interactions (Liu et al., 2014; Nowakowski and Tinoco, 1997). The high throughput sequencing data from Kobori and Yokobayashi (Kobori and Yokobayashi, 2016) contains a library of mutants including 144 single, 10,152 double, 240,967 triple and 272,690 other (>3) mutated variants. This is 100% coverage of all possible single and double mutations.

**Figure 3.**
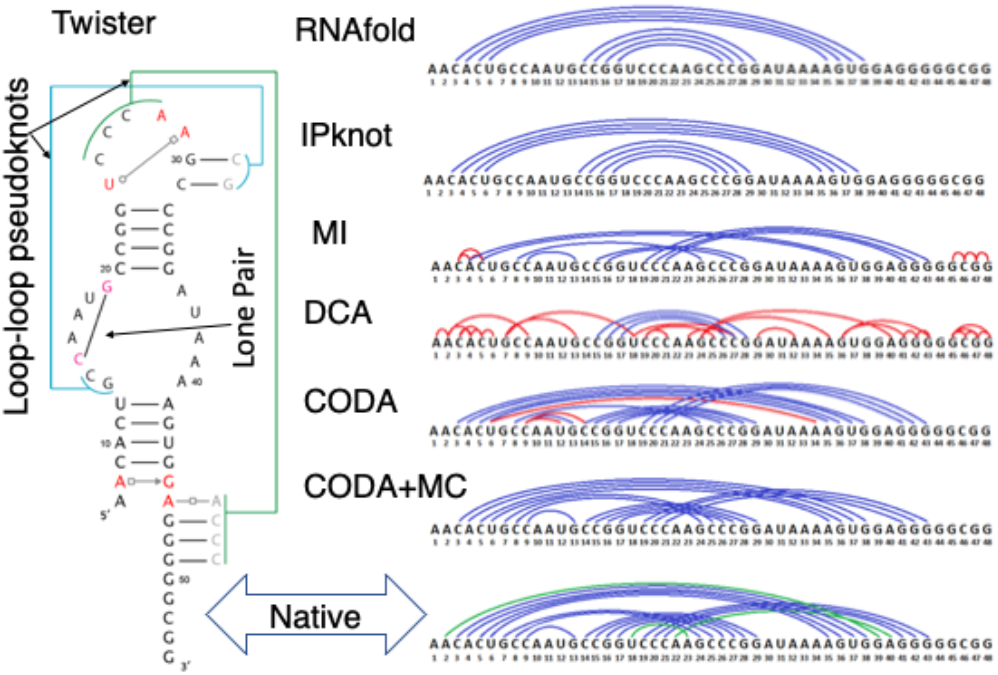
The base-pairing structure of the twister ribozyme along with base pairs predicted by secondary structure predictors RNAfold and IPknot and inferred from its deep mutation data by mutual information (MI), direct coupling analysis (DCA), covariation-induced deviation of activity (CODA), and coupling of CODA and Monte Carlo simulated annealing (CODA+MC). In the right panel, native Watson-Crick and noncanonical base pairs are shown in blue and green, respectively. False positive predictions are shown in red.

Table 1 compares the performance of various methods for the twister ribozyme. The overall trend is the same as CPEB3 ribozyme. CODA increases sensitivity by 23% and precision by 21% over MI, which is substantially better than DCA in this particular case. CODA has the highest MCC value (0.79), compared to MI (0.56), DCA (0.19), and RNAfold and IPknot (0.68). The base-pairing plot (Figure 3) shows that CODA yields overall correct topology with lone pair and pseudoknots but misses 4 base pairs with three false positive predictions. These false positives are associated with an obvious error of one base paired with more than a single base. Supplementary Figure S2 further compares the precision-recall curve given by MI, DCA, and CODA. CODA achieves 50% improvement over the next best MI in the area under the precision-recall curve. It has zero false positives (100% precision) for the first 7 base pairs (41% sensitivity), indicating that these seven base pairs can be identified without any ambiguity, compared to only 2 by MI. We noted, however, that unlike the case of CPEB3 ribozyme, all methods failed to detect three noncanonical base pairs. The density plot for the base-pairing map (Supplementary Figure S3) and score distributions (Supplementary Figure S4) confirm that the greater separation of bases unpaired and paired is the reason for high sensitivity and precision of CODA, relative to MI and DCA. More importantly, CODA+MC determines all 14 canonical base pairs at 100% precision, including lone and non-nested base pairs. Unlike CPEB3 ribozyme, all methods missed three noncanonical base pairs (A2G39, A22A40, and Hoogsteen base pair U18A22).

## DISCUSSION

Base pairing is responsible for stabilizing overall structural fold of RNA structures and the key for understanding functional mechanisms. A full base-pairing structure is more than secondary structure because it contains tertiary contacts such as lone base pairs and kissing loop pseudoknots. This work shows that these challenging tertiary contacts can be inferred from deep mutation data by analyzing covariation-induced deviation of double-mutation activity from the mutation-independent model (CODA). The combination of CODA with a Monte Carlo simulated annealing leads to detection of all including lone WC pairs for twister ribozyme at 100% precision and all WC pairs plus three non-WC pairs for CPEB3 ribozyme at 96.3% precision, despite that CPEB3 ribozyme has only a small library of <50,000 single-to-triple mutants and the mutation data of twister ribozyme is limited to the large fragment from self-cleavage. Moreover, both have two pseudoknots associated with tertiary interactions.

The CODA analysis developed here is significantly more powerful than mutual information and direct coupling analysis. This is reflected from 35% and 51% increases in the areas under the precision-recall curve by CODA over the next best (DCA for CPEB3 and MI for twister ribozymes), respectively. The large improvement in performance is due to an increase in the signal-to-noise ratio as a result of employing all mutation variants functional or non-functional in CODA. By comparison, MI and DCA analysis only rely on functional variants because they were originally developed to extract covariation signals from functional homologous sequences across different species.

Identification of noncanonical, lone, and non-nested base pairs is not a trivial exercise. Double-stranded long helical stems with many stacked base pairs are relatively easy to detect because they serve as core structural elements and a mutation at the center of these stems will have a large impact on the overall structural stability. Pseudoknots, on the other hand, are often involved with a few base pairs between two distant hairpin loops (loop-loop pseudoknots or kissing stem-loop (Forsdyke, 1995; Nowakowski and Tinoco, 1997)) as shown in Figures 1 and 3. These base pairs are considered as a part of tertiary contacts to help to stabilize the overall three-dimensional shape. CODA can capture one or more base pairs within a region of contiguous base pairs regardless if the pair is nested or non-nested (pseudoknots). In other words, it offers an excellent base-pairing topology for expanding into the full base-pairing structure. This is confirmed by a combination of CODA with MC simulated annealing, which significantly increases sensitivity while maintaining high precision.

A total of three base pairs were missed by the combination of CODA and Monte Carlo simulated annealing, all of which are noncanonical pairs. They are A2G39, A22A40 and Hoogsteen base pair U18A22 in twister ribozyme. The backbone orientations of these noncanonical pairs are quite different from Watson-Crick pairs so that all double mutations on these sites yield negative signals in CODA. By comparison, all three noncanonical base pairs in CPEB3 ribozyme (G6U41, C12A35, and U26U43) have positive signals in CODA and were correctly predicted by the combination of CODA and MC simulated annealing. However, one false positive C14G73 was predicted at the end of a helix in CPEB3. This site has a negative signal in CODA, but the signal is weaker than the energetic gain of adding a GC pair. The weak negative signal is due to the low coverage of double mutations, only one double mutant (A14T73) is observed at this site. A mutation library with a higher mutational coverage on GC pairs will likely lead to the full, exact base pair structure for CPEB3 ribozyme.

This study investigated two self-cleaving ribozymes as a proof of concept. Self-cleaving ribozymes are broadly distributed in genomes of different organisms from viroids to vertebrates (Roth et al., 2014; Webb et al., 2009; Weinberg et al., 2015). Understanding their structures and functions is only at the beginning (Jimenez et al., 2015a) with almost all the known ribozymes having interesting mechanistic differences (Jimenez et al., 2015b; Liu et al., 2017; Ren et al., 2017). This deep-mutation-based method can infer base-pairing structures at the single base-pair level with sensitivity and precision inaccessible to current multi-dimensional chemical probing methods.

The method, however, is not limited to self-cleaving ribozymes. Previously mutational scanning was applied to investigate tRNA (Li et al., 2016), RNA catalysis (Dhamodharan et al., 2017; Pitt and Ferre-D’Amare, 2010), RNA-protein interactions (Buenrostro et al., 2014; Tome et al., 2014), and RNA-RNA interactions (Auyeung et al., 2013; Puchta et al., 2016). The high-throughput techniques employed for function selection prior to sequencing includes *in vitro* affinity/activity selection (Buenrostro et al., 2014; Pitt and Ferre-D’Amare, 2010; Smyth et al., 2015; Tome et al., 2014) and *in vivo* assay according to cell growth (Li et al., 2016; Puchta et al., 2016) and fluorescence intensity (Trachman et al., 2017). The success of these ingenious *in vitro* and *in vivo* techniques for function selections indicates the wide applicability of deep mutational scanning in determining *in vitro* and *in vivo* base-pairing structures of many types of RNAs beyond self-cleaving ribozymes.

We demonstrated that the CODA analysis is capable of detecting nested and non-nested base pairs as well as some noncanonical base pairs even from a small mutant library of <5×10^4^ mutants for 81-nucleotide CPEB3 ribozyme. This suggests that the method should be applicable to RNAs with 800 bases or less for an achievable library of 5×10^6^ mutants (or 600 bases due to the current read length limit of the next-generation sequencing platforms). However, it is important to note that combining CODA with MC simulated annealing can fold the complex base-pairing pattern of the twister fragment correctly. By comparison, the secondary structure predictors RNAfold and IPknot have to use the whole sequence to achieve a reasonable prediction. For example, MCC=0.22 if using the twister fragment sequence for RNAfold, compared to 0.68 when using the whole twister sequence (Table 1). Solving fragment base-pairing structures is important for those RNAs that are too long to achieve a significant coverage of possible double mutations or too long for available sequencing techniques (>600 bases).

In addition to its own importance, the accurate, full base-pairing structure obtained from deep mutational scanning should benefit high-quality 3D structure modeling. This is because a full base-pairing structure contains not only secondary structure motif but also important tertiary contacts of lone, noncanonical, and non-nested base pairs. Thus, it can serve as a preformed, quasi-three-dimensional frame for correctly folding into the right RNA tertiary structure. Its usefulness for 3D structure modeling has been demonstrated in RNA Puzzles (blind RNA structure prediction) even with the data not yet at the single-base-pair level (Miao et al., 2017).

Moreover, the CODA analysis should be a powerful tool for analyzing deep mutation data of proteins. This transferability is reflected from our direct application of MI and DCA analysis programs to RNAs although they were originally designed for proteins (Feinauer et al., 2014; Morcos et al., 2011). Hydrogen-bonded base-pairing interactions in RNA structures are the dominant interaction with the strongest covariation signals. As a result, what emerges from the CODA analysis will be the base-base contact map in term of base pairing. On the other hand, hydrophobic packing with complementary shapes and sizes of side chains is the dominant driving force for protein folding (Dill et al., 2008). Thus, CODA analysis will lead to residue-residue contact maps based on distance proximity of amino acid residues. Because an accurate determination of protein contact maps has been demonstrated to yield high-resolution protein structure (Ovchinnikov et al., 2017), the results reported here suggest the potential of protein structural inference from deep mutational scanning (Fowler and Fields, 2014).

## Methods & Materials

### Deep mutational scanning of self-cleaving CPEB3 ribozyme

The overall procedure for deep mutational scanning of CPEB3 ribozyme is shown in Figure 1. All oligonucleotides were purchased from IDT. The sequences of oligonucleotides used in this experiment are listed in Supplementary Table S1.

Firstly, a mutation library was constructed by randomly mutating the wild type sequence of CPEB3 ribozyme with several rounds of epPCR (Cadwell and Joyce, 1992) using primer T7prom and M13F (Supplementary Table S1). The amplification product of each round was purified, quantified, and diluted for use as the DNA template in the next round of epPCR. PCR products after several rounds of epPCR were barcoded using primer Bar_F and Bar_R (Supplementary Table S1) by a low-cycle PCR (cycle number=3). The purified barcoding product was then loaded to a denaturing 8% PAGE with 8 M urea for separation. The target band containing barcoded DNAs was excised and recovered by passive elution in crush-soak buffer (10mM Tris-HCl, pH 7.5, 200mM NaCl, 5mM EDTA) overnight at 37°C followed by ethanol precipitation. The recovered DNAs were dissolved in water and then quantified by absorbance and qPCR. Approximately 10^5^ barcoded DNA molecules were amplified by T7prom and M13F primers (Table S1) in order to produce enough DNAs for the downstream steps.

Then, the mutant DNA library was transcribed *in vitro* in a 30 μl reaction system containing 5 pmol of the dsDNA template, 2mM NTPs, 1X RNAPol Reaction Buffer (New England Biolabs), 1 U/μl Murine RNase Inhibitor (New England Biolabs), and 5 U/μl T7 RNA polymerase (New England Biolabs) for 3 hours at 37 °C. Template DNA was removed by adding 58 μl nuclease-free water, 10 μl of 10X DNase I Buffer (New England Biolabs), and 2 μl of DNase I (2 U/μl, New England Biolabs) to 30 μl RNA product, and incubated at 37°C for 15 minutes. The active ribozyme mutants were simultaneously cleaved during *in vitro* transcription and DNase I treatment. 2 μl of 0.5 M EDTA (to a final concentration of 10 mM) was added to stop both ribozyme and DNase I activities. The transcribed RNAs were purified using RNA Clean & Concentrator-5 kit (Zymo Research) and quantified by absorbance.

Afterward, approximately 10 pmol purified RNAs were mixed with 2 μl of 10μM RT_m13f_adp1 (Table S1) and 1 μl of 10mM dNTP in a volume of 8 μl and heated to 65°C for 5 minutes and placed on ice. Reverse transcription was initiated by adding 4 μl of 5X ProtoScript II Buffer (New England Biolabs), 2 μl of 0.1 M DTT, 0.2 μl of Murine RNase Inhibitor (40 U/μl, New England Biolabs), 1 μl ProtoScript II RT (200 U/μl, New England Biolabs), and 2 μl 100μM template-switching oligonucleotide TSO (Supplementary Table S1) to a total volume of 20 μl. The reaction mixture was incubated at 42°C for 1 hour, then inactivated at 80°C for 5 minutes. RNA was removed by adding 1 μl 5M NaOH and heating at 95°C for 5 minutes. In Batch 2, the cDNA with template switching was used to construct RNA-seq library directly. In Batches 1 and 3, the cleaved and uncleaved cDNAs were separated by PAGE and then mixed with an equal molar ratio for RNA-seq library construction.

Next, DNA-seq libraries were constructed by extending the ribozyme mutant library with P5R1_m13f and P7R2_t7p (Supplementary Table S1) in the touch-up PCR reaction (2 cycles of PCR with the annealing temperature at 53°C, followed by 8 cycles of PCR with the annealing temperature at 67°C). Similarly, the RNA-seq libraries were constructed by extending the purified cDNA after template switching reaction with P5R1_adp1 and P7R2_adp2 (Supplementary Table S1).

Lastly, the DNA-seq and RNA-seq libraries were sequenced on an Illumina HiSeq X sequencer with 25% PhiX control by Novogene Technology Co., Ltd. The reads were first assessed using FastQC (https://www.bioinformatics.babraham.ac.uk/projects/fastqc/). Raw paired-end reads were merged using program SeqPrep (John, J.S., 2011. https://github.com/jstjohn) to generate high-quality full-length reads. For DNA-seq results, reads with a correct match to primers Bar_F and Bar_R were extracted to generate the projection map from barcodes to variants. For multiple variant sequences mapped to one barcode, the partial sequence downstream to the cleavage site was employed as the additional marker for tracking different mutants. If multiple mapping was still an issue, only the consensus sequence with the frequency larger than 50% was used. For RNA-seq results, reads with a correct match to primer Bar_R were extracted. The barcode together with partial sequences downstream to the cleavage site was used to map the full-length ribozyme sequence. Then, we counted the cleaved and uncleaved read numbers for each variant to calculate relative activity (*RA*), using the equation given by *RA(var)* = *N*_*cleaved*_*(var)N*_*total*_*(wt)*/*N*_*total*_*(var)N*_*cleaved*_*(wt)*. We performed three batches of experiments. The DNA and RNA sequencing results and the number of mutations in CPEB3 ribozyme for these batches were summarized in Supplementary Tables S2 and S3, respectively. Three different batches with different conditions were used to diversify our mutants with different mutation rates (See Supplementary Figure S5 for the mutation rates at each position). To obtain a higher coverage of mutation variants, we merged the RA results together. Average RA was used if a variant appeared in different batches. Because the coverage for double mutations in CPEB3 ribozyme was low (41.0%), we increased the coverage by including triple mutants in our CODA analysis as follows. For a triple mutant XYZ, we calculate *RA*(XY) = *RA*(XYZ)/*RA*(Z) when *RA*(Z) is larger than 0.5. Here we have assumed that Z mutation does not covariate with XY mutations. If *RA*(XY) is covered by the double mutation data, we employed *RA*(XY)=0.7 *RA*^(2)^(XY)+0.3 *RA*^(3)^(XY) where *RA*^(2)^(XY) and *RA*^(3)^(XY) are from double and triple mutations, respectively. In this way, all triple mutants were included as double mutants in CODA analysis. This extension increased the double mutation coverage from 41.0% to 61.3% (Supplementary Table S5). For consistency, the same approach was used to include triple mutations in twister ribozyme.

### Covariation-induced deviation of activity (CODA)

The overview of the method is shown in Figure 1C. Specific details are described below.

#### The independent-mutation model

The basic assumption of CODA analysis is that the effect of two mutations on enzymatic activity is independent if they are not in close contact. This assumption is an approximation in the case that long-range interactions or allosteric effects are important. Under this approximation, the *RA* of double mutants can be modeled (predicted) by the *RA*s of two single mutants. However, if two bases form a base pair, the *RA* of their double mutation to another complementary pair will be higher than two independent single mutations that are disruptive to the wild-type pair. In other words, we can detect base pairs by examining COvariation-induced Deviation of the Activity (CODA) of a double mutant from the independent-mutation model. We constructed this model by using an SVR regression of the *RA* of double-mutation variants against *RA*s of their corresponding single-mutation variants for those likely unpaired bases. This was done by organizing *RA* data into a three-dimensional vector of *RA*_obs_(bi), *RA*_obs_ (bj), and *RA*_obs_(bi, bj), which are *RA*s of two single mutation variants and the double mutation variant, respectively. Then, the data points of likely unpaired bases (with *RA*_obs_(bi, bj) smaller than either *RA*_obs_ (bi) or *RA*_obs_ (bj)) are used to generate the SVR model to fit *RA*_obs_(bi, bj) as a function of *RA*_obs_ (bi) and *RA*_obs_ (bj). This SVR model is used to predict the activity for any double mutations as we have 100% coverage for single mutations of twister and CPEB3 ribozymes. Comparison between observed and predicted *RA* values is shown in Figure 1D for CPEB3 ribozyme and Supplementary Figure S6 for a twister ribozyme. The SVR regression model was generated using the python package scikit-learn with the parameter C=2×10^3^ and gamma=2.0 to optimize the regression. The model is based on deep mutation data only without requiring the knowledge of base pairs.

#### CODA calculation

The CODA score is defined as the difference between the actual *RA* value (*RA*_obs_) and the predicted *RA* (*RA*_pred_) from the above regression model normalized by a shifted predicted *RA* value. *CODA*=*(RA*_*obs*_−*RA*_*pred*_)/*(RA*_*pred*_+*0.2)*. A shift (0.2) is used to avoid the small value of *RA*_*pred*_. The results do not depend on this value from 0.1 to 0.5.

#### CODA distribution

We expected that there are two populations in the distribution of CODA scores for double mutations from an arbitrary pair to the canonical AU or GC pairs: one centered near 0 for structurally unpaired bases (two mutations are independent) and one centered at a higher value (outliers) for potential base pairs as illustrated in Figure 1D and Supplementary Figure S6. We defined outliers as three standard deviations away from the baseline and then fits those outliers to a Gaussian distribution for base pairs and the remaining CODA scores were used to fit another Gaussian distribution to represent unpaired bases. For CPEB3 ribozyme, the distributions for paired and unpaired have mean values at 1.168 and −0.011 and deviation at 0.751 and 0.214, respectively, with the overall base-pair probability (P(paired)) at 0.015. For twister ribozyme, the distributions for paired and unpaired have mean values at 1.541 and −0.098 and deviation at 0.505 and 0.189, respectively, with the overall pair probability P(paired)=0.019. To tolerate the inaccuracy of division of paired and unpaired scores, the deviation in the fitted Gaussian distribution for presumed base pairs is two times of the observed value. We have tested several different ways to generate these two distributions. They only yielded minor changes in distributions without affecting the final outcome of the calculations. Again, there is no specific assumption about which two bases are paired or not paired.

#### Pairing score by the Bayes classifier

The pairing score of a double mutation (*Ps*) is calculated by a naïve Bayes two-state classifier with with

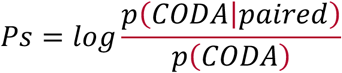

with
*p*(*CODA*) = *p*(*CODA*|*paired*) * *p*(*paired*) + *p*(*CODA*|*unpaired*) * (1 − *p*(*paired*)). This classifier is based on a statistical analysis of deep mutation data only without requiring the knowledge of specific base pairs.

### Secondary structure prediction by Monte Carlo simulated annealing

We built a simple secondary structure predictor to examine if it can be used to fill the gap in the region of contiguous base pairs with weak covariation signals. We defined the energy score as follows.

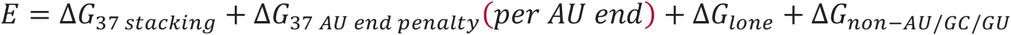

where the stacking energy Δ*G*_37 *stacking*_ and the penalty of helical AU end Δ*G*_37 *AU end penalty*_(*per AU end*) were obtained from Tuner 2004 experimental data (Xia et al., 1998). Δ*G*_*lone*_ and Δ*G*_*non-AU/GC/GU*_ are the penalties for a lone pair and for non-AU, GC, or GU (wobble) pairs, respectively. To make the simplest model, we used a single value for both Δ*G*_*single*_ and Δ*G*_*non-AU/GC/GU*_ penalties. We found the low penalty value by trials and errors to ensure this simple secondary structure prediction not to yield any lone or non-AU/GC/GU pair. In each Monte Carlo step, a random existing pair was destroyed, or a new pair was generated. The move is accepted according to the Metropolis criterion. At each temperature, we performed 500,000 Metropolis steps. The initial temperature was set at 10 and decreased by a factor of 0.95 at each round until temperature reached 0.1. The results were stable when the single adjustable penalty parameter varied between 2-5 kcal/mol.

After the above energy function was defined, we carried out the CODA-guided base-pairing prediction by using *E-Ps* without using any weight factor because they were in the same order of magnitude. The simulated annealing procedure was the same as above for using MC alone.

### Covariation analysis by mutual information and direct coupling analysis

#### Defining functional sequences for MI and DCA by an RA cutoff

Functional variants with high ribozyme activity were used as the input sequences for covariation analysis by MI and DCA. We have tested different RA cutoffs (0.2, 0.3, …, 0.9) for defining functional sequences. The result was not sensitive to *RA* between 0.5 and 0.8 (See Supplementary Figure S7). As a result, we mainly reported the result with *RA* cut off at 0.5 (Table 1).

#### Mutual information

Mutual information (MI) was calculated by the following equation:

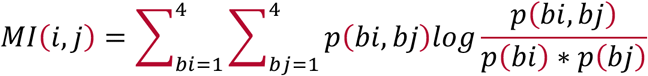

where *i* and *j* are two different base positions, *bi* and *bj* are base types (A, U, G, C) at position *i* and *j*, *p(bi)* and *p(bj)* are the frequency of the base type at each position, *p(bi,bj)* is the frequency of the base pair at these two positions.

#### Direct coupling analysis

DCA was calculated by gplmDCA method using default parameters (Feinauer et al., 2014). The program was downloaded from https://github.com/mskwark/gplmDCA.

### Performance measurement

For binary classification of bases paired and unpaired, we measured the performance of different methods by Matthews correlation coefficient (MCC), sensitivity (recall), and precision. MCC, sensitivity, and precision are defined by

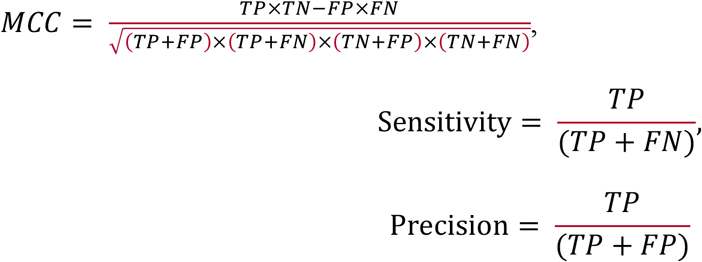

where *TP*, *TN*, *FP*, and *FN* represent True Positive (base pairs), True Negative (two bases unpaired), False Positive (two bases unpaired but predicted as paired) and False Negative (two bases paired but predicted as unpaired), respectively. Sensitivity (or recall) is a measure of the fraction of correctly predicted base pairs in all known base pairs (i.e. the percent of the coverage of all native base pairs). Precision is a measure of the fraction of correctly predicted base pairs in all predicted base pairs. RNA bases unpaired are about 50 times more than base pairs. For such an unbalanced system, MCC is a balanced measure (Boughorbel et al., 2017). MCC is determined by a threshold that separates paired from unpaired. The maximum value of MCC along with sensitivity and precision at the same cut off are reported here for the comparison between different methods. We also employed the precision-recall curve rather than the receiver operator characteristic curve because our main interest is the minor class of positive samples (Boughorbel et al., 2017) (in this case, base pairs).

#### RNAfold and IPknot

Secondary structure predictions using RNAfold and IPknot were calculated through their web servers at http://rna.tbi.univie.ac.at/cgi-bin/RNAWebSuite/RNAfold.cgi and https://rtips.dna.bio.keio.ac.jp/ipknot/, respectively.

#### Self-cleaving twister ribozyme

Twister ribozyme data and processing perl scripts were provided by Dr. Yokobayashi (Kobori and Yokobayashi, 2016). 86,459,834 uncleaved and 30,016,488 cleaved reads were extracted from the raw data. Relative activity of each variant was calculated by the following equation: *RA(var)* = *N*_*cleaved*_*(var)N*_*total*_*(wt)*/*N*_*total*_*(var)N*_*cleaved*_*(wt)*.

## Acknowledgment

We are indebted to Dr. Shungo Kobori and Professor Yohei Yokobayashi for providing us the deep mutation data of the twister ribozyme. Z.Z. gratefully acknowledges the award of Griffith University and University of Chinese Academy of Sciences (GU-UCAS Joint Doctoral Program) Postgraduate Scholarships. This work was supported in part by Australia Research Council DP180102060 and by National Health and Medical Research Council (1121629) of Australia to Y.Z. We would like to acknowledge the use of the High-Performance Computing Cluster ‘Gowonda’ to complete this research. This research/project has also been undertaken with the aid of the research cloud resources provided by the Queensland Cyber Infrastructure Foundation (QCIF) and Australian Research Data Commons (ARDC).

## Author Contribution

JZ and JW designed experimental studies and wrote the manuscript; ZZ and JZ conducted experimental studies and wrote the manuscript; PX developed computational methods and wrote the manuscript. TZ assisted in computational analysis. YZ conceived of the study, participated in the initial design, assisted in data analysis, and drafted the manuscript; and all authors read, contributed to the discussion, and approved the final manuscript.

