## Supplement file for "Accurate inference of the full base-pairing structure of RNA by deep mutational scanning and covariation-induced deviation of activity"

Table S1. Oligonucleotides used in this study.

| Name | Sequence (5' - 3') | Notes |
|---|---|---|
| CPEB3_wt | TAATACGACTCACTATAGGGATAACAGGGGGCCACAGCAGAA GCGTTCACGTCGCAGCCCCTGTCAGATTCTGGTGAATCTGCGA ATTCTGCTGTATATCTCATTTG AGTTACTCAGGATTACTGGCCGTCGTTTTAC | DNA template. |
| CPEB3_C57U | TAATACGACTCACTATAGGGATAACAGGGGGCCACAGCAGAA GCGTTCACGTCGCAGCCCCTGTCAGATTCTGGTGAATCTGTGA ATTCTGCTGTATATCTCATTTG AGTTACTCAGGATTACTGGCCGTCGTTTTAC | DNA template. |
| M13F | GTAAAACGACGGCCAGT | PCR primer. |
| T7prom | TAATACGACTCACTATAGGG | PCR primer. |
| RT_m13f_adp1 | CCCTACACGACGCTCTTCCGATCTGTAAAACGACGGCCAGT | Reverse transcription primer. |
| TSO | CTCGGCATTCCTGCTGAACCGCTCTTCCGATCTrGrGrG | Template switching oligo. |
| Bar_F | TAATACGACTCACTATAGGGA | PCR primer for adding barcode. |
| Bar_R | GTAAAACGACGGCCAGTNNNNNNNNNNNNNNNAATCCTGAG TAACTCAAAT | PCR primer for adding barcode. |
| P5R1_m13f | AATGATACGGCGACCACCGAGATCTACACTCTTTCCCTACAC GACGCTCTTCCGATCTGTAAAACGACGGCCAGT | PCR primer for DNA-seq library preparation. |
| P7R2_t7p | CAAGCAGAAGACGGCATACGAGATCGGTCTCGGCATTCCTGC TGAACCGCTCTTCCGATCTTAATACGACTCACTATAGG | PCR primer for DNA-seq library preparation. |
| P5R1_adp1 | AATGATACGGCGACCACCGAGATCTACACTCTTTCCCTACAC GACGCTCTTC | PCR primer for RNA-seq library preparation. |
| P7R2_adp2 | CAAGCAGAAGACGGCATACGAGATCGGTCTCGGCATTCCTGC TGAAC | PCR primer for RNA-seq library preparation. |

Table S2. Summary of deep sequencing results of CPEB3 ribozyme in three batches of experiments.

| | Number of DNA-seq reads | Number of Barcodes | Number of RNA-seq reads | Cleaved | Uncleaved |
|---|---|---|---|---|---|
| Batch 1 | 9,834,235 | 54,502 | 16,490,545 | 41.4% | 58.6% |
| Batch 2 | 28,107,628 | 89,397 | 32,061,733 | 17.8% | 82.2% |
| Batch 3 | 23,028,064 | 115,977 | 90,995,324 | 49.4% | 50.6% |

Table S3. Fractions of CPEB3 ribozyme mutants in three batches of experiments

| Number of Mutations | Batch 1 | Batch 2 | Batch 3 |
|---|---|---|---|
| 0 | 28.5% | 0.9% | 5.3% |
| 1 | 30.6% | 4.0% | 14.5% |
| 2 | 22.1% | 9.8% | 22.2% |
| 3 | 12.0% | 16.0% | 22.1% |
| 4 | 4.9% | 19.6% | 16.9% |
| 5 | 1.5% | 18.7% | 10.5% |
| 6 | 0.4% | 14.3% | 5.3% |
| 7 | 0.1% | 9.4% | 2.3% |
| 8 | 0.0% | 5.0% | 0.8% |
| 9 | 0.0% | 2.2% | 0.3% |

Table S4. Mutation rates of CPEB3 ribozyme in three batches of experiments

| Wild type | Batch 1 | Batch 2 | Batch 3 |
|---|---|---|---|
| A | 2.96% | 8.86% | 6.18% |
| U | 3.39% | 8.84% | 6.35% |
| G | 0.50% | 2.97% | 1.57% |
| C | 0.40% | 2.59% | 1.37% |
| all | 1.73% | 5.62% | 3.7% |

Table S5. The number of variants and the coverage of double mutations of CPEB3 ribozyme in three batches of experiments

| # Mutations | 1 | 2 | 3 | 4 | 5 | Double mutation coverage1 | Double mutation coverage2 |
|---|---|---|---|---|---|---|---|
| Batch 1 | 239 | 4781 | 4897 | 1771 | 476 | 16.4% | 25.1% |
| Batch 2 | 240 | 5161 | 11534 | 12165 | 9651 | 17.7% | 36.7% |
| Batch 3 | 243 | 9566 | 22409 | 15933 | 8498 | 32.8% | 52.9% |
| Merged | 243 | 11968 | 36214 | 29657 | 18567 | 41.0% | 61.3% |

- Double mutation coverage1: only use information from double mutants
- Double mutation coverage2: use information from both double mutants and triple mutants, for triple mutant XYZ, $RA(XY) = RA(XYZ)/RA(Z)$ when $RA(Z)>0.5$.

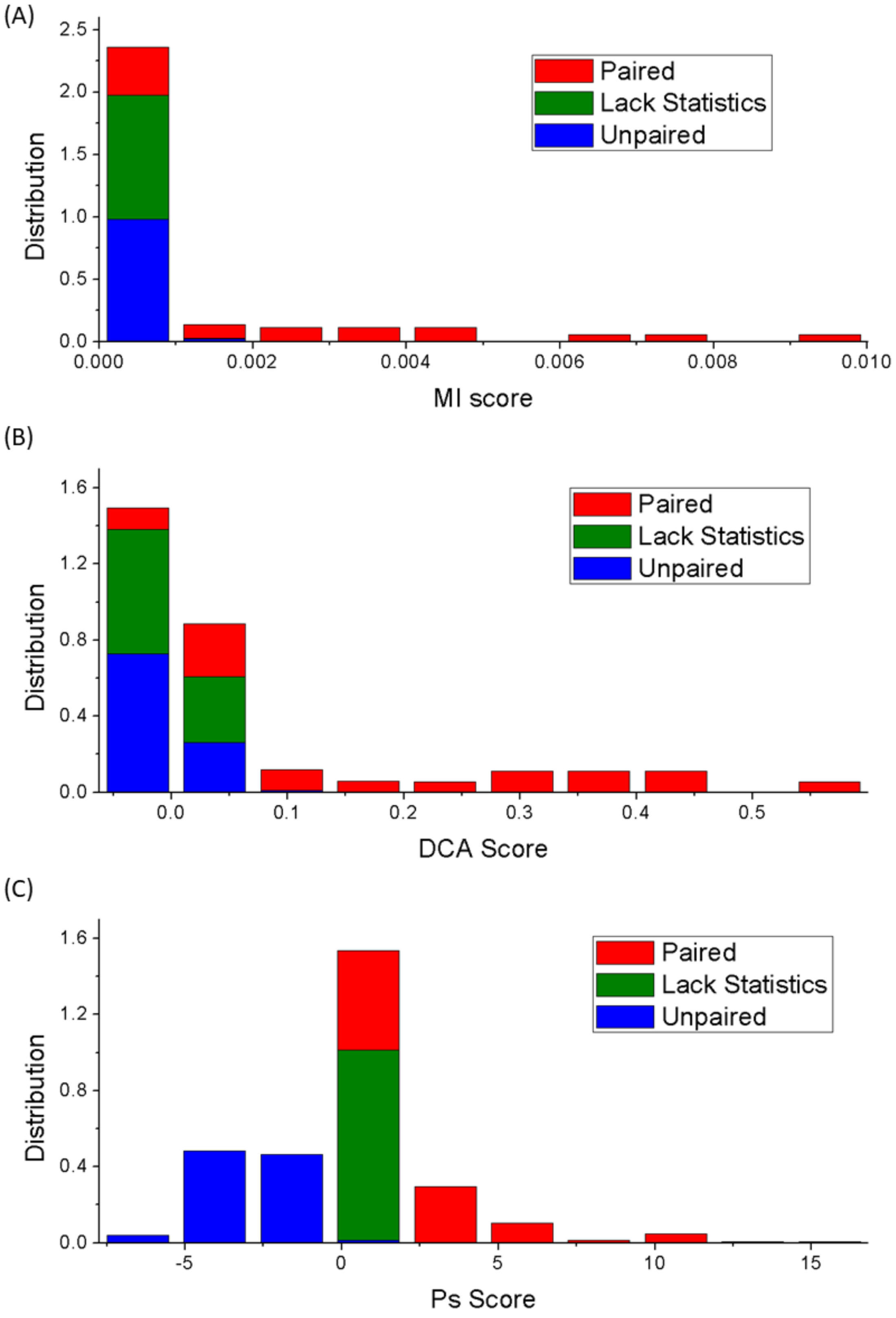


Figure S1. Distributions of MI (A), DCA (B), and Pairing Ps (CODA) (C) scores for bases paired, unpaired, and lack of statistics in CPEB3 ribozyme, respectively.

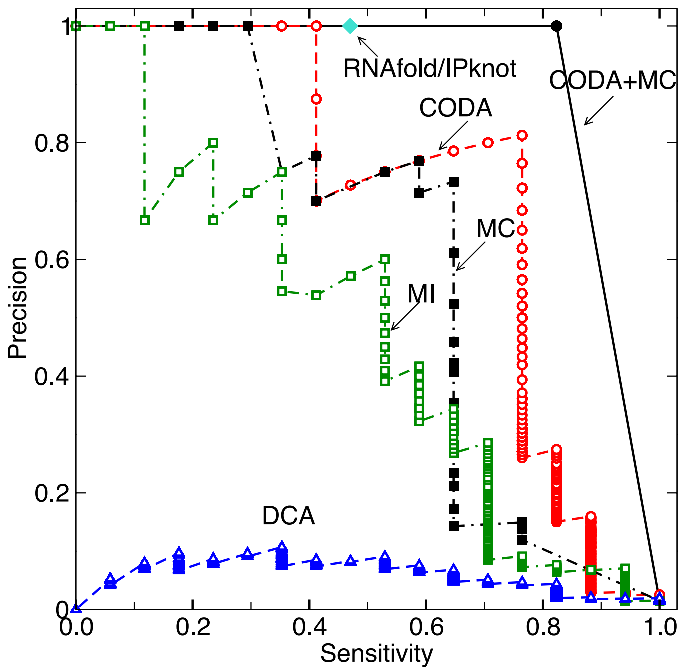


Figure S2. Precision (fraction of correct base pairs in predicted base pairs) versus sensitivity (coverage of known base pairs) by mutual information (MI), direct coupling analysis (DCA), covariation-induced deviation of activity (CODA), Monte Carlo (MC) simulated annealing, and coupling of CODA and MC, using the deep mutation data from twister. Identical results from secondary structure predictor (RNAfold and IPknot) are also shown as a point (diamond).

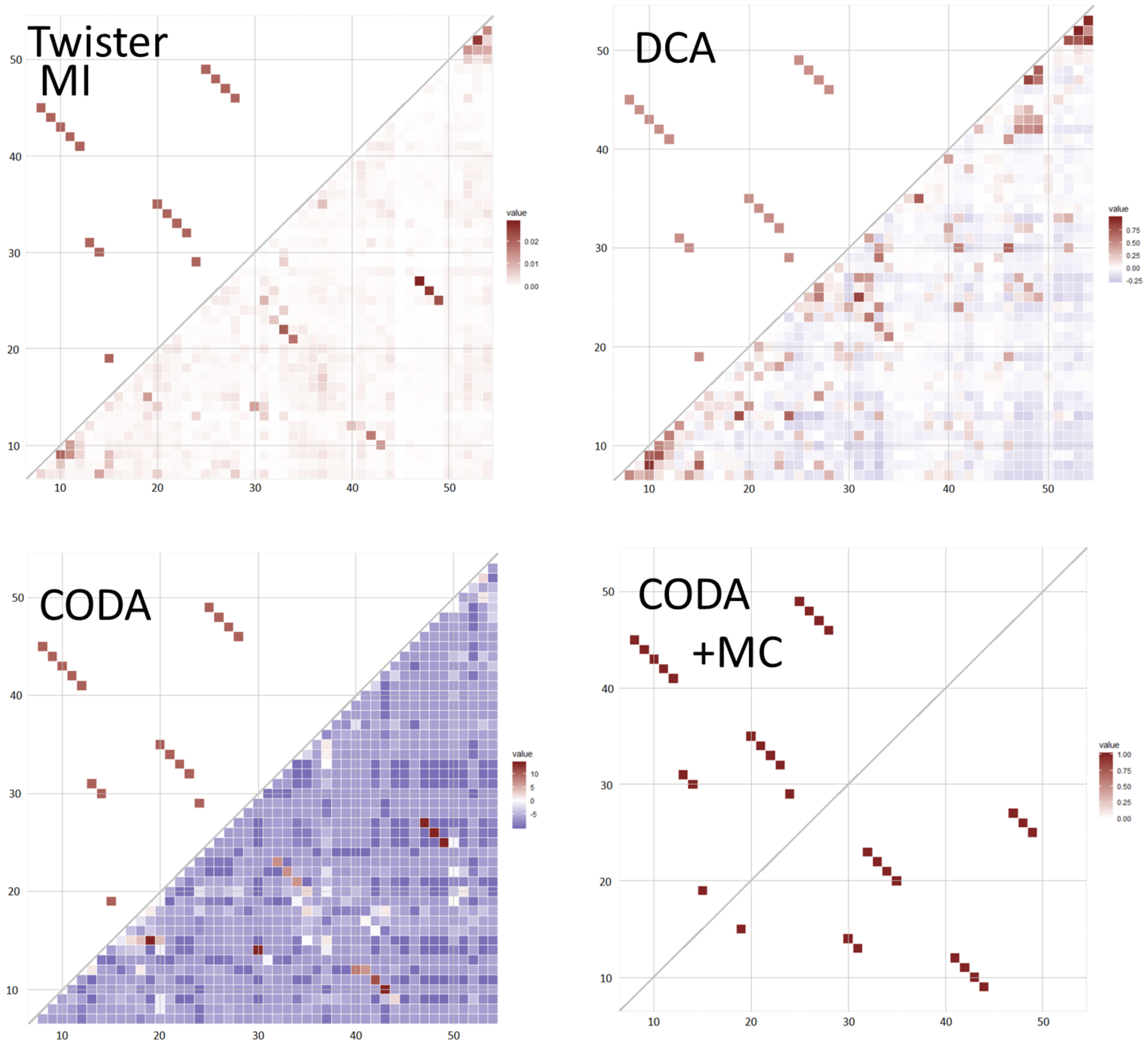


Figure S3. The comparison between native base-pairing map (upper triangle) of twister ribozyme and the map inferred from deep mutation data (lower triangle) of twister ribozyme by MI, DCA, CODA, and CODA+MC as labeled.

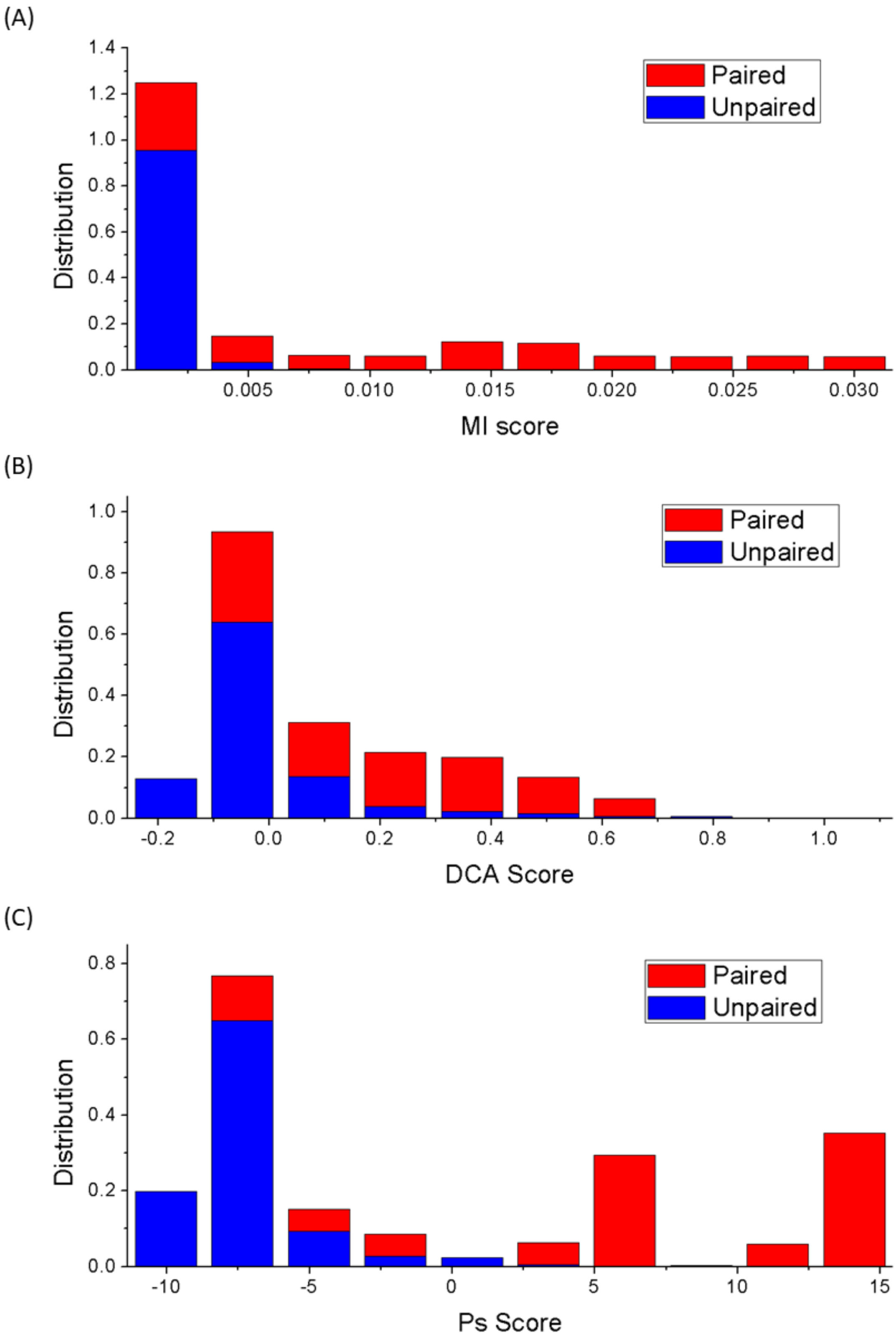


Figure S4. Distribution of MI (A), DCA (B), and Paring Ps (CODA) (C) scores for bases paired and unpaired in twister ribozyme, respectively.

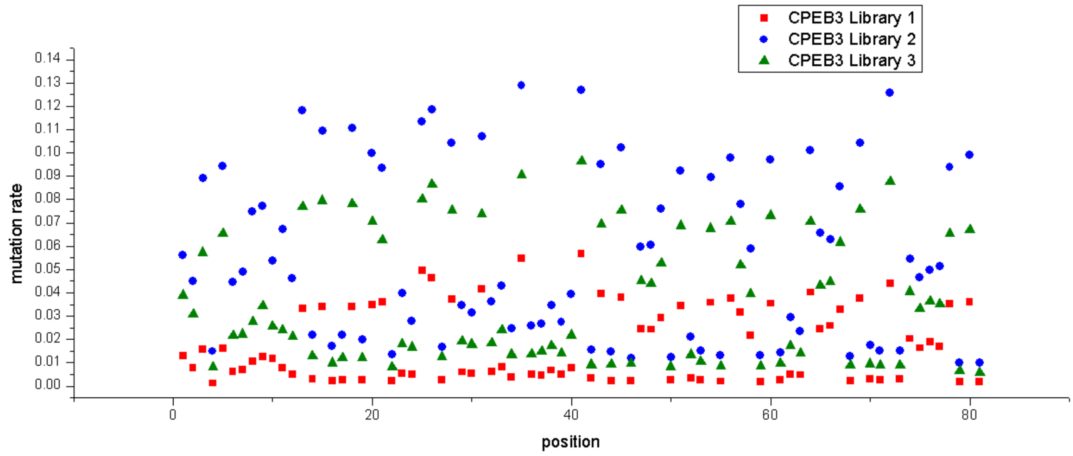


Figure S5. Mutation rates of three batches at each nucleotide position of CPEB3 ribozyme.

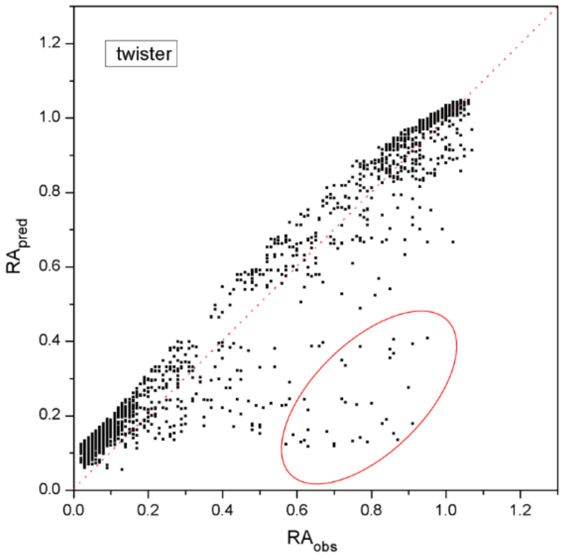


Figure S6. Observed relative activities (*RA*) of the double mutants of twister ribozyme are compared to predicted *RA* by the regression model, the red circle indicates the outliers with significant covariation-induced deviations of activity from the regression model.

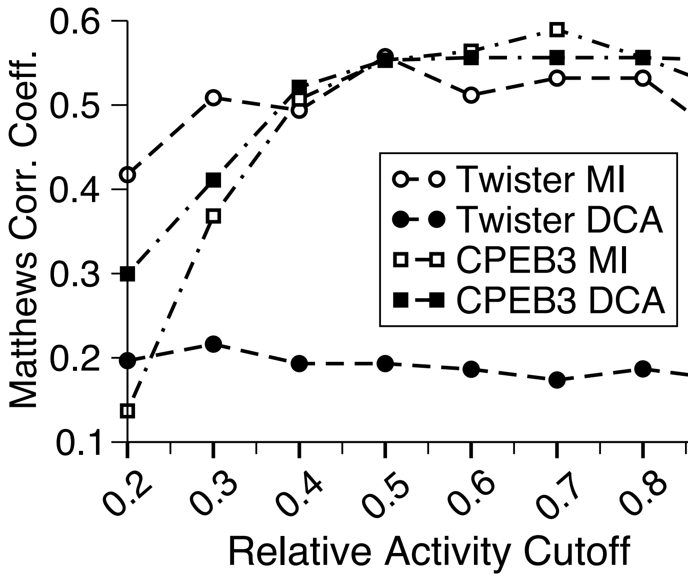


Figure S7. The performance of mutual information and direct coupling analysis in term of Matthews correlation coefficient as a function of the cut-off for defining functional ribozymes (twister and CPEB3 ribozymes as labeled).
